## Supplementary File 2 - qRT-PCR conditions for "Evidence for pre-climacteric activation of AOX transcription during cold-induced conditioning to ripen in European pear (*Pyrus communis* L.)"

**Supplemental File 2.** Quantitative RT-PCR reaction conditions and thermal profile.

*Quantitative reverse transcription PCR reagent*: BioRad Laboratories iTaq Universal SYBR Green Supermix with ROX reference dye.

*Quantitative reverse transcription PCR instrument:* Stratagene MX3005P

*Reaction components (single reaction):*

- 10.0 μl iTaq Universal SYBR Green Supermix with ROX.
- 1.0 μl forward primer at 5uM.
- 1.0 μl reverse primer at 5uM.
- 8 μl water and 100ng 1st strand cDNA.

*Reaction thermal profile*:

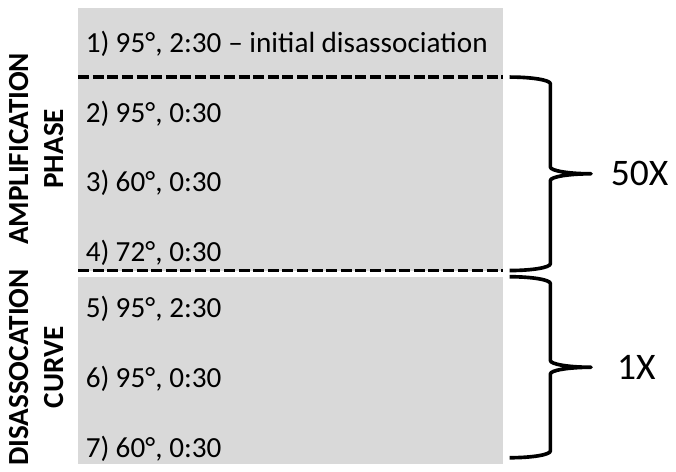
