## Supplementary figures and images for "Evidence for pre-climacteric activation of AOX transcription during cold-induced conditioning to ripen in European pear (*Pyrus communis* L.)"

### Supplementary File 5 - NMDS1_ordination by treatment - phenology overlay

#### Slide 1
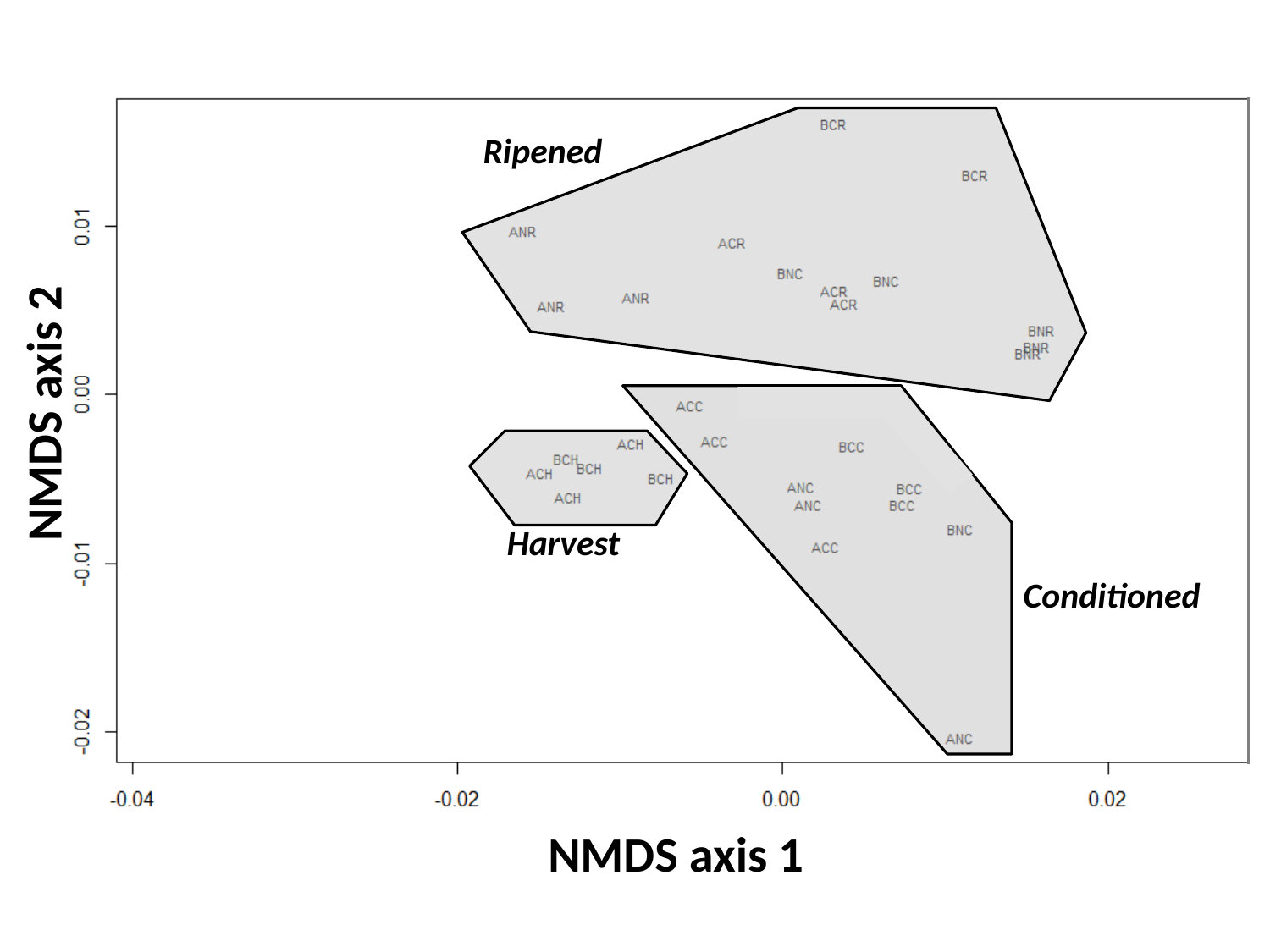

Ripened
NMDS axis 2
Harvest
Conditioned
NMDS axis 1

### Supplementary File 11 - Fig3_RawRoutput_CentroidHullPlot

#### Slide 1
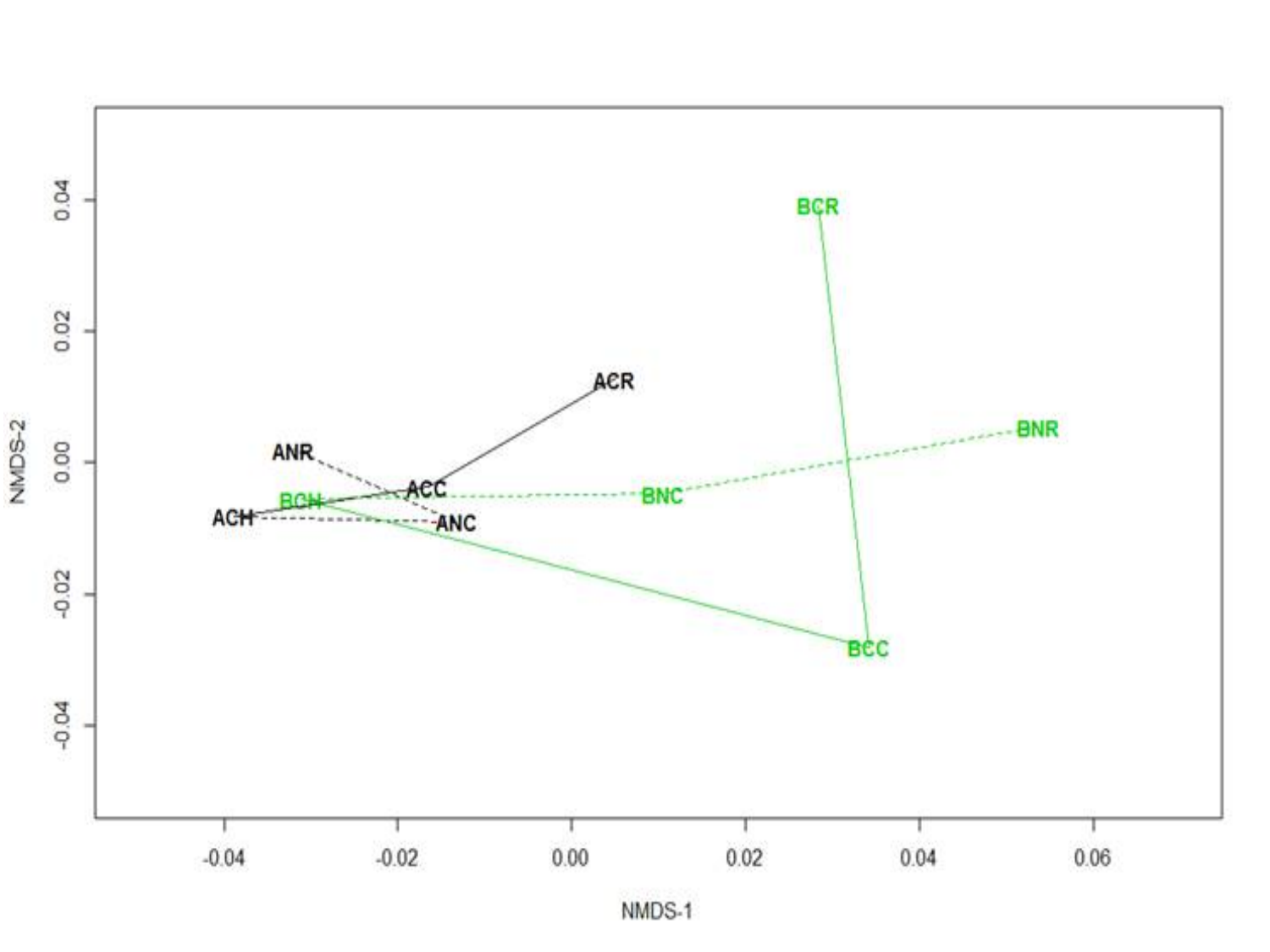

NMDS axis 2
NMDS axis 1
0.04
0.04
0.00
-0.02
-0.04
-0.04
-0.02
0.00
0.02
0.04

### Supplementary File 12 - Fig4_RawRoutput_RayBiplot

#### Slide 1
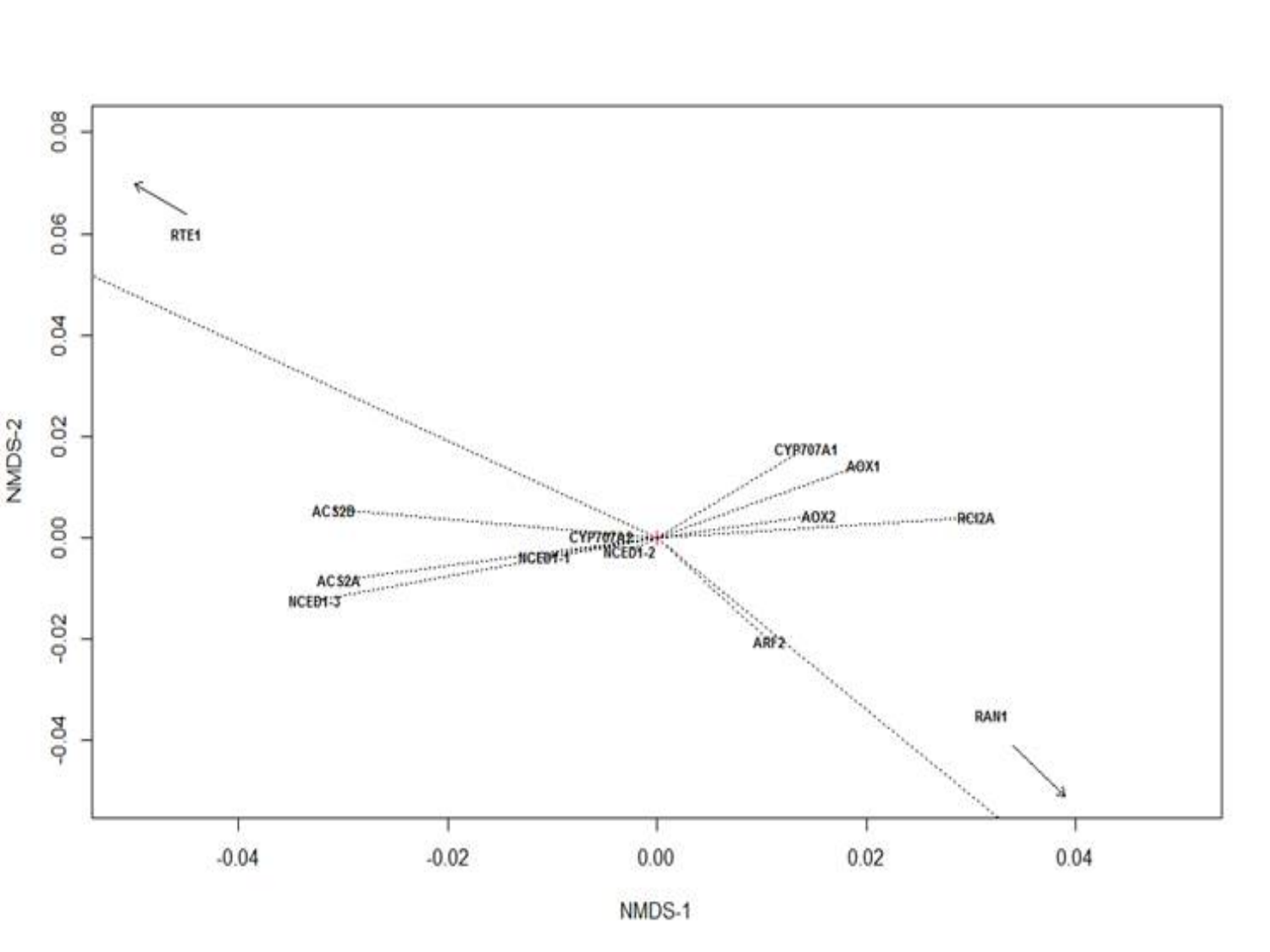

### Supplementary File 13 - RawMorpheus

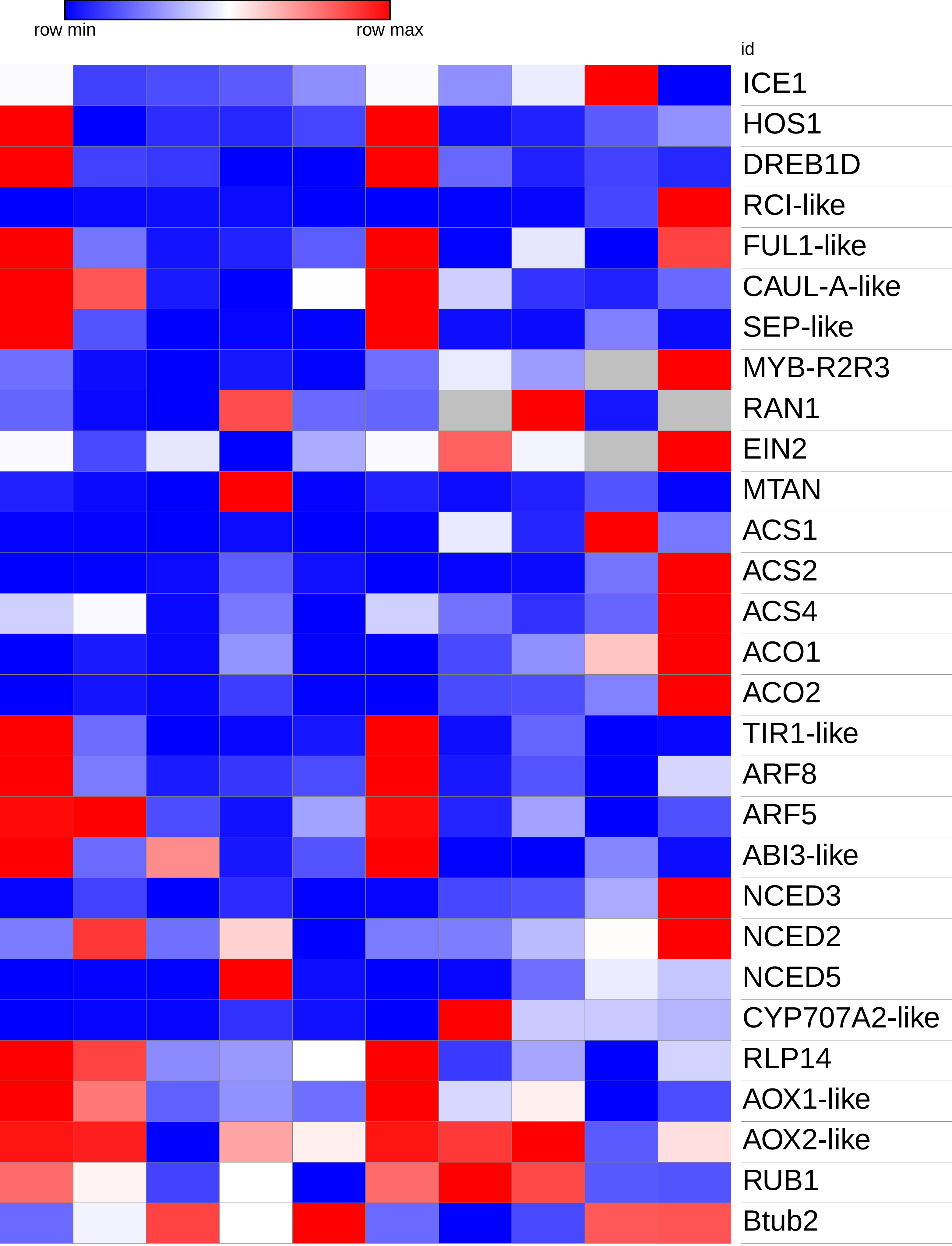
