## Supplementary File 8 - Stress plots NMDS1 and 2 for "Evidence for pre-climacteric activation of AOX transcription during cold-induced conditioning to ripen in European pear (*Pyrus communis* L.)"

**Supplemental File 8**. (A, top) Stress plots of initial (circles) and second (triangles) NMDS ordination procedures. Both instances produced a final stress coefficient of nearly 0.20 after 20 iterations. *-One gene target (VAS1) replicated (technical replicates) 3 times only. (B, bottom) Dissimilarity plot from ordination, with non-metric correlation and linear fit.

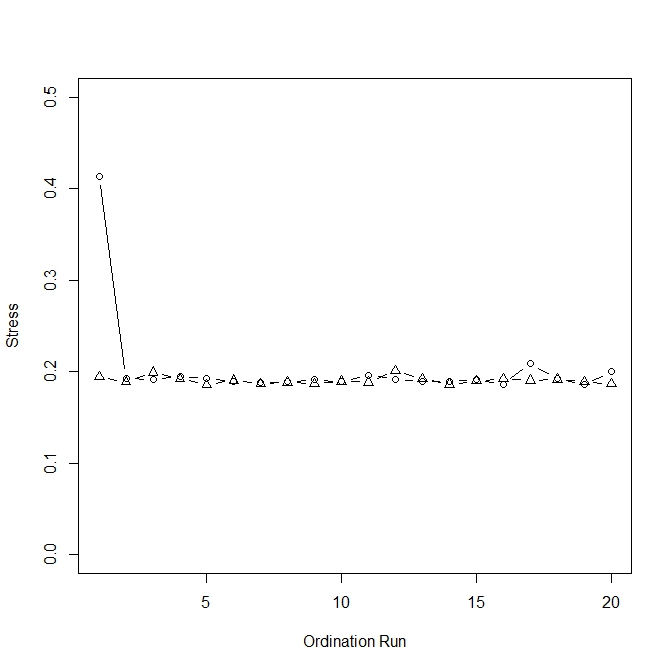

NMDS1 – 92 gene targets, 1 technical replicate

NMDS2 – 36 gene targets, 4 technical replicates*

NMDS

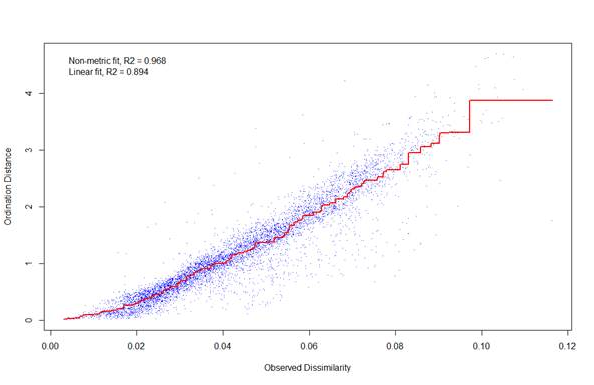
