## Supplementary File 10 - RadialDistance_NMDS1 and 2 for "Evidence for pre-climacteric activation of AOX transcription during cold-induced conditioning to ripen in European pear (*Pyrus communis* L.)"

Linear distance of genes’ variability (of expression data) in ordination space of initial NMDS procedure from initial technical quantitative RT-PCR replicate reactions among 90 genes (top) and among final 28 genes with (total of) 4 technical quantitative RT-PCR replicates (below). NMDS axis 1 strongly discriminates pear cultivar. NMDS axis 2 strongly discriminates fruits’ relative ripeness.

| **NMDS axis 1** | **NMDS axis 2** | **Gene** | **Distance** | **CAHP Number** |
| --- | --- | --- | --- | --- |
| -0.012576445 | 0.047004888 | CYP707A2 | 0.048658 | 160 |
| -0.045241431 | 0.002587101 | ACO1 | 0.045315 | 8 |
| -0.040076332 | 0.005372213 | ACO2 | 0.040435 | 1 |
| -0.030643612 | -0.014825848 | LPH14 | 0.034042 | 15 |
| -0.025060163 | 0.01488136 | ACS2 | 0.029146 | 14 |
| 0.018809424 | 0.021608828 | ARF2 | 0.028648 | 81 |
| 0.012061003 | 0.025570255 | RTE1 | 0.028272 | 51 |
| -0.027544581 | -0.00354082 | ACS1 | 0.027771 | 12 |
| -0.021248456 | 0.015210187 | NCED5 | 0.026131 | 159 |
| 0.001994823 | -0.018058458 | GLP1 | 0.018168 | 142 |
| -0.015939822 | 0.003068384 | NCED2 | 0.016232 | 158 |
| 0.013150569 | 0.007863117 | MADS-Rin-like2 | 0.015322 | 23 |
| 0.000106244 | 0.013940259 | TIR1-like | 0.013941 | 84 |
| -0.011958145 | -0.007048232 | ACS4 | 0.013881 | 17 |
| -0.005235097 | -0.012836832 | ACS3 | 0.013863 | 18 |
| 0.012181851 | 0.00598182 | ABI5 | 0.013571 | 67 |
| 0.013122827 | 0.00345279 | MYB-R2R3 | 0.013569 | 77 |
| -0.012400685 | 0.004187189 | NCED3 | 0.013089 | 157 |
| 0.01239551 | 0.002950398 | HOS1 | 0.012742 | 30 |
| -0.009348204 | -0.007691981 | CAL-A | 0.012106 | 168 |
| 0.00857949 | -0.008186749 | ABA2 | 0.011859 | 61 |
| 0.010661167 | -0.004614023 | ERF109-like | 0.011617 | 64 |
| 0.009306238 | 0.00574053 | SEP-like | 0.010934 | 24 |
| 0.008380875 | -0.006290509 | ABA1 | 0.010479 | 60 |
| -0.00411843 | -0.009371488 | RAN1 | 0.010237 | 52 |
| 0.00895562 | 0.003616075 | ICE1 | 0.009658 | 46 |
| 0.007056636 | 0.006116478 | Cullin-1 | 0.009338 | 47 |
| -0.00218599 | -0.008936934 | MTAN | 0.0092 | 45 |
| 0.009142884 | -0.000673577 | ARF8 | 0.009168 | 82 |
| 0.009008724 | -0.001130903 | FUL1/MADS3 | 0.009079 | 169 |
| -0.004646063 | 0.007340283 | CYP707A1 | 0.008687 | 162 |
| 0.006979608 | 0.004249644 | DREB1D | 0.008172 | 35 |
| 0.007825643 | -0.000498264 | ARF5 | 0.007841 | 80 |
| 0.005478824 | 0.005255358 | CTR1 | 0.007592 | 4 |
| -0.002433605 | -0.006820471 | ERS1A | 0.007242 | 27 |
| 0.00511446 | 0.005090221 | AOX1 | 0.007216 | 164 |
| 0.007045859 | -0.000675302 | ERF-LP1 | 0.007078 | 59 |
| 0.006916686 | 0.00108892 | MADS-Rin like1 | 0.007002 | 22 |
| -0.000507812 | -0.006865516 | ETR1A | 0.006884 | 25 |
| -0.000956817 | -0.006786637 | CDPK28 | 0.006854 | 2 |
| -0.006224026 | 0.001754006 | CBF2 | 0.006466 | 73 |
| -0.000840982 | -0.006359423 | CDPK1 | 0.006415 | 34 |
| 0.005844275 | -0.002339735 | TAG1 | 0.006295 | 151 |
| 0.00043232 | -0.006190157 | ERF1B | 0.006205 | 54 |
| 0.000934045 | -0.006075108 | VIN3-like | 0.006146 | 153 |
| -0.002439215 | -0.005539408 | HOS2 | 0.006053 | 50 |
| 0.0049775 | -0.003430174 | ERS1B | 0.006045 | 28 |
| 0.005725704 | 0.000269656 | AdoMet synthase | 0.005732 | 69 |
| 0.001504401 | -0.005344073 | AC4 | 0.005552 | 125 |
| 0.00185746 | -0.005005038 | ER-Auxin binding protein | 0.005339 | 165 |
| -0.000614747 | -0.005204567 | ERF3B | 0.005241 | 58 |
| 0.000841799 | -0.0050711 | EZA1 | 0.00514 | 129 |
| 0.004654606 | -0.00158237 | IAA14 | 0.004916 | 100 |
| -0.003978834 | -0.002845605 | GH3-1 | 0.004892 | 137 |
| 0.002493444 | -0.003840942 | AMI1 | 0.004579 | 87 |
| 0.002886075 | -0.003441263 | ERF3A | 0.004491 | 57 |
| 0.004010863 | 0.00156314 | ABP1 | 0.004305 | 103 |
| 0.003866638 | -0.001705233 | PP2C-like | 0.004226 | 62 |
| 0.004064816 | -0.000809178 | VRN1 | 0.004145 | 134 |
| 0.003606535 | 0.001995638 | SKP1 | 0.004122 | 90 |
| 0.003307763 | 0.002264281 | IAA3.1 | 0.004009 | 91 |
| 0.003945701 | 0.000146638 | OPCL1 | 0.003948 | 127 |
| 0.001661053 | -0.00353605 | XBAT32 | 0.003907 | 31 |
| 0.002024445 | 0.003234752 | DRIP2 | 0.003816 | 76 |
| 0.002721805 | -0.002249086 | ARF4 | 0.003531 | 81 |
| 0.000966144 | -0.003204563 | ERF2B | 0.003347 | 56 |
| 0.003151279 | 0.000676059 | ERF60 | 0.003223 | 74 |
| 0.002303931 | -0.002236639 | ICE2 | 0.003211 | 33 |
| 0.002849158 | 0.000572733 | NADP malic enzyme | 0.002906 | 110 |
| 0.000971177 | -0.002444211 | EOL1 | 0.00263 | 32 |
| 0.001740107 | -0.001719246 | ETR1B | 0.002446 | 26 |
| -0.001036562 | -0.002110549 | EBF1 | 0.002351 | 21 |
| -0.00225541 | -0.00029576 | FRY1 | 0.002275 | 49 |
| -0.000129807 | -0.001873332 | EIL2 | 0.001878 | 20 |
| -0.001327284 | 0.00129858 | DREB2B | 0.001857 | 75 |
| 0.001810851 | -0.000321428 | PIN5 | 0.001839 | 136 |
| 0.000788717 | -0.001588883 | VRN2 | 0.001774 | 135 |
| -0.000124623 | -0.001758468 | NIT1 | 0.001763 | 98 |
| 0.001371751 | 0.000928203 | LEA14 | 0.001656 | 88 |
| -0.000289959 | -0.001447799 | CLM | 0.001477 | 71 |
| 0.000305831 | -0.001432166 | ERF2A | 0.001464 | 55 |
| 0.0000612 | -0.001411327 | AFP-like | 0.001413 | 68 |
| 0.001369804 | 0.000332625 | SR1 | 0.00141 | 150 |
| 0.000704277 | -0.001163653 | ERF1A | 0.00136 | 53 |
| 0.00038794 | -0.000923556 | EIN5 | 0.001002 | 97 |
| 0.000887304 | 0.000459711 | SBP1 | 0.000999 | 147 |
| -0.000224685 | 0.000897004 | MTK1 | 0.000925 | 70 |
| -0.000358103 | 0.000762925 | EIN2 | 0.000843 | 10 |
| 0.000491772 | 0.00032423 | RUB1 | 0.000589 | 78 |
